# Altered PVN-to-CA2 hippocampal oxytocin pathway and reduced number of oxytocin-receptor expressing astrocytes in heart failure rats

**DOI:** 10.1101/2022.03.17.484603

**Authors:** Ferdinand Althammer, Ranjan K. Roy, Arthur Lefevre, Rami Najjar, Kai Schoenig, Dusan Bartsch, Marina Eliava, Rafaela Feresin, Elizabeth Hammock, Anne Z. Murphy, Alexandre Charlet, Valery Grinevich, Javier E. Stern

**Author notes:** Corresponding author: **Javier E. Stern**, M.D. Ph.D., Center for Neuroinflammation and Cardiometabolic Diseases, Georgia State University, Atlanta, GA 30302-5030 United States.

## Abstract

Oxytocinergic actions within the hippocampal CA2 are important for neuromodulation, memory processing and social recognition. However, the source of the OTergic innervation, the cellular targets expressing the OT receptors (OTRs) and whether the PVN-to-CA2 OTergic system is altered during heart failure (HF), a condition recently associated with cognitive and mood decline, remains unknown. Using immunohistochemistry along with retrograde monosynaptic tracing, RNAscope and a novel OTR-Cre rat line, we show that the PVN (but not the supraoptic nucleus) is an important source of OTergic innervation to the CA2. These OTergic fibers were found in many instances in close apposition to OTR expressing cells within the CA2. Interestingly, while only a small proportion of neurons were found to express OTRs (∼15%), this expression was much more abundant in CA2 astrocytes (∼40%), an even higher proportion that was recently reported for astrocytes in the central amygdala. Using an established ischemic rat heart failure (HF) model, we found that HF resulted in robust changes in the PVN-to-CA2 OTergic system, both at the source and target levels. Within the PVN, we found an increased OT immunoreactivity, along with a diminished OTR expression in PVN neurons. Within the CA2 of HF rats, we observed a blunted OTergic innervation, along with a diminished OTR expression, which appeared to be restricted to CA2 astrocytes. Taken together, our studies highlight astrocytes as key cellular targets mediating OTergic PVN inputs to the CA2 hippocampal region. Moreover, provides the first evidence for an altered PVN-to-CA2 OTergic system in HF rats, which could potentially contribute to previously reported cognitive and mood impairments in this animal model.

## Introduction

The hippocampus is one of the most critical brain structures involved in spatiotemporal memory and navigation (1-4) and hippocampal pathological alterations or damage have been associated with memory loss and cognitive decline (5-8).

The hypothalamic neuropeptide oxytocin (OT) mediates a plethora of prosocial, emotional and memory-related functions in the brain (9-13) and OT-mediated activation of hippocampal neurons has been demonstrated to fine-tune hippocampal microcircuits (14) by enhancing spike transmission in fast-spiking interneurons (15) and transforming the firing mode of hippocampal CA2 neurons (16). In fact, OT receptor (OTR)-expressing neurons can be found in various subregions of the mouse dorsal hippocampus, with the strongest presence of OTRs in the CA2 region (17). A recently generated transgenic OXTR-Venus mouse knockin line also displayed sparse populations of OTR-positive cells in CA1, CA2 and CA3 hippocampal layers (18). Moreover, OT fibers have been previously shown in the hippocampus (16, 19) and it was suggested that the paraventricular nucleus of the hypothalamus (PVN) is the primary source for this innervation (16, 19). The current view is that axonally-released OT directly acts on hippocampal interneurons and the potential involvement of other cell types in the OTergic modulation of hippocampal activity is currently unknown. Interestingly, astrocytes were recently described as key elements of OTergic modulation in the local amygdala CeL→ CeM microcircuit (20). Thus, we aimed here to determine if other brain regions, namely the hippocampus, harbor OTR-expressing astrocytes as well.

We recently described cognitive decline, impaired memory and changes in hippocampal gene expression as a direct result of heart failure (HF) in rats (21). To probe whether OTergic PVN→ CA2 connectivity or OTR expression in hippocampal CA2 was altered in the disease, we performed a series of experiments in sham and HF rats. Here we report that in stark contrast to an overall increased expression of OT in the PVN, we found a diminished OTergic innervation density of hippocampal CA2, as well as reduced OTR mRNA and protein in CA2. Intriguingly, we identified OTR-positive astrocytes in the hippocampal CA2 region, which – to the best of our knowledge – is the first account of OTRs in hippocampal astrocytes of rats. Unexpectedly, we also observed a drastic reduction in the number of OTR-expressing astrocytes in CA2 in HF rats, suggesting that this population of astrocytes is sensitive to cardiovascular-related, pathological alterations of OTergic signaling in the brain. No differences in the number of CA2-projecting PVN OT neurons were observed between sham and HF rats. Taken together, our results provide the first evidence supporting the existence of an OTR-positive subpopulation of hippocampal astrocytes, and build the foundation for future studies focused on addressing the functional role of OTR-expressing astrocytes in the hippocampus under healthy and disease conditions.

## Materials and methods

All performed experiments in rats were approved by the Georgia State University Institutional Animal Care and Use Committee (IACUC) and carried out in agreement with the IACUC guidelines. At all time, animals had *ad libitum* access to food and water and all efforts were made to minimize suffering and the numbers of animals used for this study.

### Animals

We used male Wistar rats (5-7 weeks old at surgery, 180-200g, Envigo, Indianapolis, IN, USA) for all experiments (total n=93). For the analysis of OTR-expressing neurons in dorsal CA2, we used the recently developed OTR-Cre knockin rats ((27), n=3). Rats were housed in cages (2 per cage) under constant temperature (22 ± 2°C) and humidity (55 ± 5%) on a 12-h light cycle (lights on: 08:00-20:00). OTR mice (*Oxtr*^*tm1*.*1knis*^) were maintained on a C57BL/6J background (30, 33) and bred at Florida State University on a 12L:12D forward cycle in wire top caging with wood chip bedding, a nestlet, and ad libitum access to food (LabDiet 5001) and water. All procedures were performed after approval by The Institutional Animal Care and Use Committee, Florida State University (protocol #1722 and #1746) in accordance with state and federal guidelines (Guide for the Care and Use of Laboratory Animals of the National Institutes of Health). Mice were bred from untimed co-housed heterozygous pairs and checked daily for litters. The first appearance of a new litter was designated post-natal day 0 (P0). On P14, mice were given an overdose of sodium pentobarbital and transcardially perfused with 0.1M phosphate buffer (pH 7.4) followed by 4% paraformaldehyde in phosphate buffer (pH 7.4). Brains were removed and post-fixed for 24 hours in 4% paraformaldehyde at 4°C followed by subsequent changes in escalating concentrations of sucrose in PBS (10, 20, 30% for 24 hours each). After genotype confirmation of mice (66), 2 OTR wild-type and 2 OTR knock-out brains (all females) were cryosectioned in the coronal plane at 40 microns in 3 series and stored at −20°C in cryoprotectant. Samples were shipped to Georgia State University and stored frozen until immunofluorescence.

### Heart failure surgery and Echocardiography

As previously described (67) HF was induced by coronary artery ligation surgery. Animals were anaesthetized using 4% isoflurane and intubated for mechanical ventilation until the end of the surgery. To exteriorize the heart, we performed a left thoracotomy. The ligation was performed on the main diagonal branch of the left anterior descending coronary artery. Animals received buprenorphine SR-LAB (0.5 mg/kg, S.C.; ZooPharm, Windsor, CO, USA) before the surgical procedure to minimize postsurgical pain. Sham animals underwent the same procedure except the occlusion of the left coronary artery. Four to five weeks after the surgery we performed transthoracic echocardiography (Vevo 3100 systems; Visual Sonics, Toronto, ON; Canada) under light isoflurane (2-3%) anesthesia to assess the ejection fraction (EF) and confirm the development of HF. We obtained the left ventricle internal diameter and the left diameter of the ventricle posterior and anterior walls in the short-axis motion imaging mode to calculate the EF. The myocardial infarct surgery typically results in a wide range of functional HF, as determined by the EF measurements. Rats with EF<40% were considered as HF (Sham 86.23±2.7%, HF 36.40±2.4%).

### Immunohistochemistry

Following pentobarbital-induced anesthesia (Euthasol, Virbac, ANADA #200-071, Fort Worth, TX, USA, Pentobarbital, 80mg/kgbw, i.p.), rats were first perfused at a speed of 20mL/min with 0.01M PBS (200mL, 4°C) through the left ventricle followed by 4% paraformaldehyde (PFA, in 0.3M PBS, 200mL, 4°C), while the right atrium was opened with an incision. Brains were post-fixated for 24 hours in 4% PFA at 4°C and transferred into a 30% sucrose solution (in 0.01M PBS) at 4°C for 3-4 days. For immunohistochemistry, 40 µm slices were cut using a Leica Cryostat (CM3050 S) and brain slices were kept in 0.01M PBS at 4°C until used for staining. Brain slices were blocked with 5% Normal Horse Serum in 0.01M PBS for 1h at room temperature. After a 15-min washing in 0.01M PBS, brain slices were incubated for 24hrs (48hrs in case of PS38) in 0.01M PBS, 0.1% Triton-X, 0.04% NaN_3_ containing different antibodies: 1:100 PS38 Neurophysin-I (gift from Harold Gainer), 1:1000 anti-glutamine synthetase (monoclonal mouse, Merck Milipore, MAB 302, clone GS-6), anti-GFAP (goat polyclonal, abcam, ab53554) and anti-synaptophysin (anti-rabbit, abcam, ab32127, 1:1.000) at room temperature. Following 15-min washing in 0.01M PBS, sections were incubated in 0.01M PBS, 0.1% Triton-X, 0.04% NaN_3_ with 1:500 Alexa Fluor 488/594-conjugated donkey anti-rabbit/goat/mouse (Jackson ImmunoResearch, 711-585-152, 705-585-147, 715-545-151) for 4 hours at RT. Brain slices were washed again for 15 mins in 0.01M PBS and mounted using antifade mounting medium (Vectashield with DAPI, H-1200B).

### RNAScope

RNAScope reagents were purchased from acdbio (Multiplex v2 RNAScope kit). Nuclease-free water and PBS were purchased from Fisher Scientific. Brains were processed as described under ***Immunohistochemistry*** using nuclease-free PBS, water, PBS and sucrose. We followed the manufacturer protocol with a few modifications: 1) Immediately after cryosectioning, slices were washed in nuclease-free PBS to remove sucrose and OCT compound. 2) Hydrogen peroxide treatment was performed with free floating sections prior to slice mounting. 3) Sections were mounted in nuclease-free PBS at room temperature. 4) Pretreatment with Protease III was performed for 20 minutes at room temperature. 5) No target retrieval step was perfomed.

### Confocal microscopy and 3D IMARIS analysis

Confocal images were obtained using a Zeiss LSM 780 confocal microscope (1024×1024 pixel, 16-bit depth, pixel size 0.63-micron, zoom 0.7). For the three-dimensional reconstruction 40µm-thick z-stacks were acquired using 1µm-steps. Three-dimensional reconstruction of astrocytes or axons was performed as previously described (22). Image processing, three-dimensional reconstruction and data analysis was performed in a blind manner in regards to the experimental conditions.

### Analysis of OTR intensity, density and counting of cells and fibers

Density and intensity of OTR immunosignal were assessed in Fiji (NIH) using the threshold paradigm. Average pixel intensity was calculated using the Analyze<Measure function using collapsed z-stack images. For density analyses, raw images were opened and threshold was set individually for each image to achieve a near-perfect overlap of raw immunosignal and pixels used for quantifications. Optical density was then calculated using the calibrated images. For quantifications we used at least 6-8 sections for each animal and brain region, if not indicated otherwise. Cells/neurons were counted manually using the cell counter plugin in Fiji. We used 8 sections containing dorsal CA2 per animal and sections were selected according to the indicated Bregma levels. OTergic fibers were counted manually in Fiji, while we defined a continuous line longer 2µm as a separate fiber.

### Reverse transcription polymerase chain reaction (RT-PCR) and quantitative real time PCR (qPCR)

RNA extraction and isolation were performed using the miRNAeasy Mini kit (Qiagen, Cat. No. 217004) and the QIAzol Lysis Reagent (Qiagen, Mat. No. 1023537). 200µm-thick tissue sections were made in cryostat (−20°C, Leica, CM3050S) and punches from the PVN (8-12 punches per animal) and DH (10-14 punches per animal) were collected and kept in dry ice until the RNA extraction procedure. RNA concentration was measured using NanoDrop One (Thermo Scientific) and was in the range of 240 – 370 ng/µl prior to cDNA synthesis. cDNA synthesis was performed using the iScript™ gDNA Clear cDNA Synthesis Kit (BIO RAD, cat. no. 1725035) and the SimpliAmp Thermal Cycler (applied biosystems, Thermo Fisher Scientific) according to the manufacturer protocol. qPCR was conducted using the following 10x QuantiTect primers (diluted in 1.1 mL TE pH 8.0, final concentration: 200nM) purchased from Qiagen: OTR (QT100379036) and β-Actin (QT00193473), used as the reference gene). All individual qPCR reactions (brain region, primer and condition) were triplicated and averages for further statistical analysis.

### Stereotaxic injection of viral vectors

Injection of viral vectors into the rat brain was performed in analogy to (19). If not indicated otherwise, all hypothalamic nuclei were injected bilaterally using the following coordinates: SON (M-L ± 1.6mm, A-P −1.4mm, D-V −9.0mm) and PVN (M-L ± 0.3mm, A-P −1.8mm, D-V −8.0mm). Injections into dorsal CA2 were performed using the following coordinates: (M-L ± 3.5mm, A-P −3.0mm, D-V −3.5mm and M-L ± 3.5mm, A-P −3.5mm, D-V −3.5mm). Point of origin for the coordinates was Bregma and the Z level difference of Bregma and Lambda did not exceed 0.1mm (68). Injection volume per injection site was 300 nl, while all viruses used were in the range of 10^12^ −10^13^ genomic copies per ml.

### Brain-region specific western blots using tissue punches

Brain punches were homogenized in radioimmunoprecipitation assay (RIPA) buffer supplemented with phosphatase inhibitor cocktail 1 and 2 and protease inhibitor cocktail (Sigma Aldrich, Saint Louis, MO) followed by centrifugation at 16,000 g for 20 min at 4°C. Protein from supernatant was quantified and normalized via the DC protein assay kit (BioRad Laboratories, Hercules, CA). In preparation for sodium dodecyl sulfate (SDS) polyacrylamide gel electrophoresis, 40 μg of samples 3 were mixed with 2× laemmli buffer + 5% 2-mercaptoethanol (BioRad Laboratories, Hercules, CA). Samples were then briefly vortexed and centrifuged, then heated for 10 min at 70 °C in a dry heating block and loaded onto a polyacrylamide gel for electrophoresis. Following electrophoresis, gels were transferred to a polyvinylidene difluoride (PVDF) membrane using a Trans-Blot Turbo Transfer System (Bio-Rad Laboratories, Hercules, CA). Membranes were then blocked in TBS-T (50 m mol/L Tris, 150 m mol/L NaCl, 0.2% Tween-20, pH 7.4) + 5% non-fat dry milk (NFDM) and washed in TBS-T (3 × 5 min). Membranes were incubated overnight in 4 °C with mouse monoclonal antibody (dilution 1:1000 in TBS-T + 5% BSA) oxytocin receptor (Cat #: sc-515809) from Santa Cruz Biotechnology (Dallas, TX) and rabbit monoclonal antibody GAPDH (Cat #: 2118) from Cell Signaling (Danvers, MA). Membranes were then washed in TBS-T (3 × 5 min) and probed with secondary antibodies: mouse (Cat #: 7076S; Cell Signaling Danvers, MA) or rabbit (Cat #: 7074; Cell Signaling Danvers, MA) (1:5,000 dilution) in TBS-T + 5% NFDM for 1 h at room temperature. After three 5-min washes with TBS-T, chemiluminescent imaging of target proteins was performed with Immobilon Forte Western HRP Substrate (EMD Millpore, Billerica, MA) using the ChemiDoc Imaging Systems (BioRad Laboratories, Hercules, CA). Band density of oxytocin receptor was quantified using Image Lab 6.0 (BioRad Laboratories, Hercules, CA) and normalized to GAPDH.

### Statistical analysis

All statistical analyses were performed using GraphPad Prism 8 (GraphPad Software, California, USA). The significance of differences was determined using Student’s t test or one-sample t-test, as indicated in the respective figure legends. Chi-Square tests were used to compare differences in the incidence of OTR+ cells between sham and HF rats. Results are expressed as mean ± standard error of the mean (SEM). Results were considered statistically significant if p<0.05 and are presented as * for p<0.05, ** for p<0.01 and *** for p<0.0001 in the respective Figures.

## Results

### HF-induced changes in OT immunoreactivity and OTR protein levels in the PVN

To investigate potential HF-induced changes in OT neurons within the PVN, we three-dimensionally reconstructed OT-immunoreactive cells using the Imaris software (22). (**Figure 1A-E**). As summarized in Figures 1A, B, we observed an overall increased OT immunoreactivity in the PVN of HF rats, as shown by an increase in OT fluorescent signal intensity. While we did not find differences in the total number of OT neurons (**Figure 1C, D)**, we did observe an increase in the total number of OT-positive voxels (**Figure 1E**), which is also indicative of increased OT expression and possibly somatodendritic swelling. This was further corroborated by assessing soma area of OT neurons, which was significantly increased in HF rats (**Figure 1F**). Next, to determine whether the expression of oxytocin receptors (OTR) was also altered in the PVN of HF rats, we performed western blot for OTR using PVN tissue punches (**Figure 1G, H**). Notably, we found that OTR protein expression in the PVN of HF rats was downregulated by almost 50%, potentially indicating a robust adaption to the previously reported increase in OTergic activity in HF rats (23).

**Figure 1.**
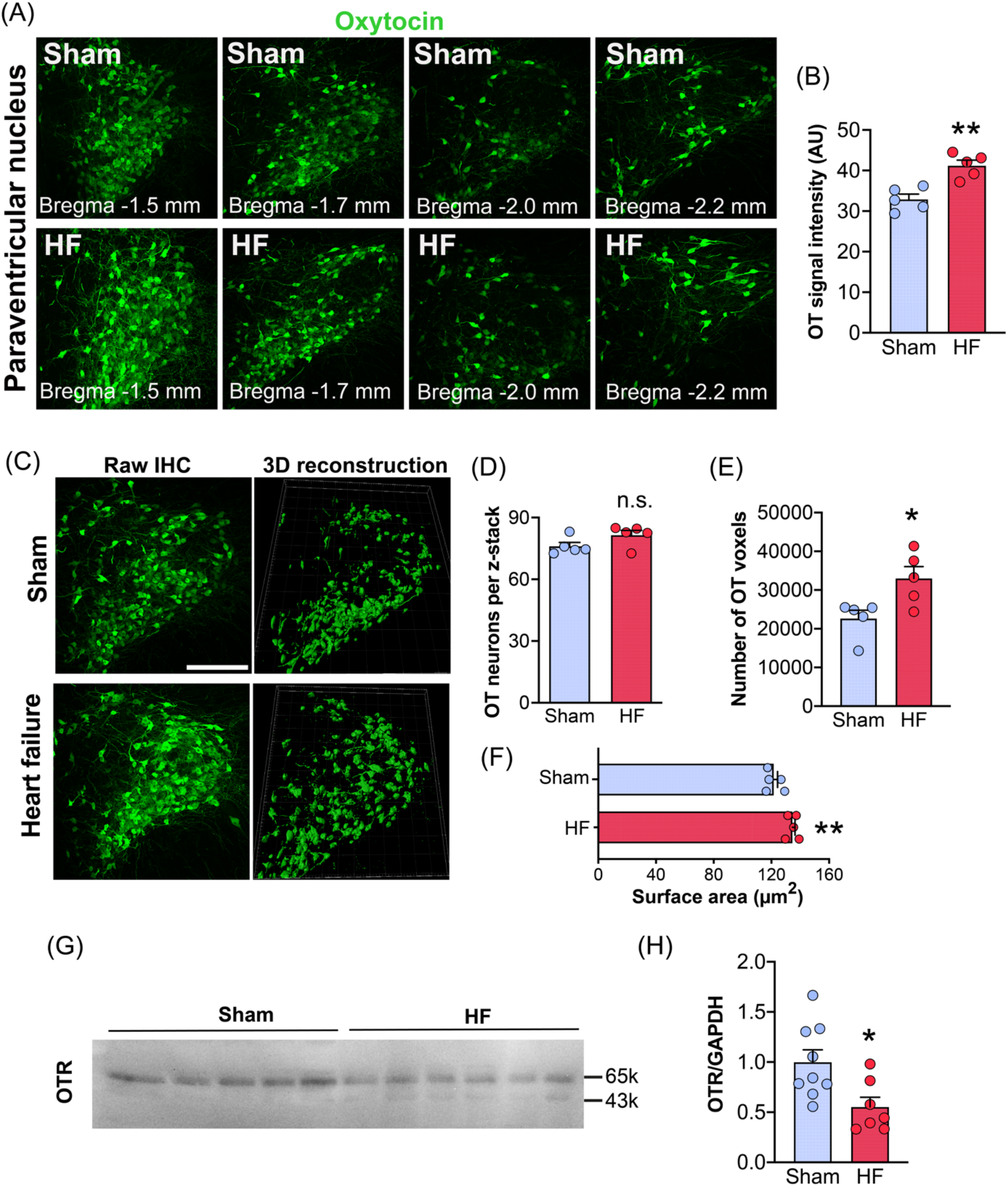
(A) Confocal images of PVN OT neurons of HF and sham rats and various different rostro-caudal levels (Bregma: −1.5mm, −1.7mm, −2.0mm and −2.2mm). (B) Increased OT immunoreactivity in PVN OT neurons of HF rats. Bar graph shows pooled data from all rostro-caudal PVN regions. (C) Confocal images (left panels) and three-dimensional reconstructions (right panels) of OT neurons within the PVN stained via PS38 OT antibody. Scale bar=250µm. (D) No difference in the total number of OT PVN neurons between sham and HF rats (n=5 per group). (E) Increased number of OT-positive voxels in HF rats. (F) OT neurons in HF rats have enlarged somata (n=5 per group). (G) Western blot images for OTR and GAPDH (control) for sham and HF rats. (H) Reduced OTR protein levels in HF rats (n=8 sham, n=6 HF). Student’s t-test, *p<0.05 and **p<0.001.

### HF does not affect the number of PVN→ CA2 projections

OTergic innervation of CA2 has been described before in mice and rats (24, 25), albeit these reports remain controversial (26). Thus, we specifically probed for the presence of OTergic fibers in the CA2 of the rat. We observed sparse OTergic innervation of CA2 (**Figure 2A**), with very little to no innervation of CA1 or CA3 (not shown), and found the highest number of OTergic fibers at Bregma - 3.6mm (**Figure 2B**). Intriguingly, we occasionally observed basket-like, OTergic puncta on CA2 neurons (**Figure 2C**), which – to the best of our knowledge – have only been described before in mice (16), but not in rats. We used three-dimensional reconstruction of these OTergic puncta and confirmed their close proximity to CA2 neurons located in the pyramidal cell layer (**Figure 2D**).

**Figure 2.**
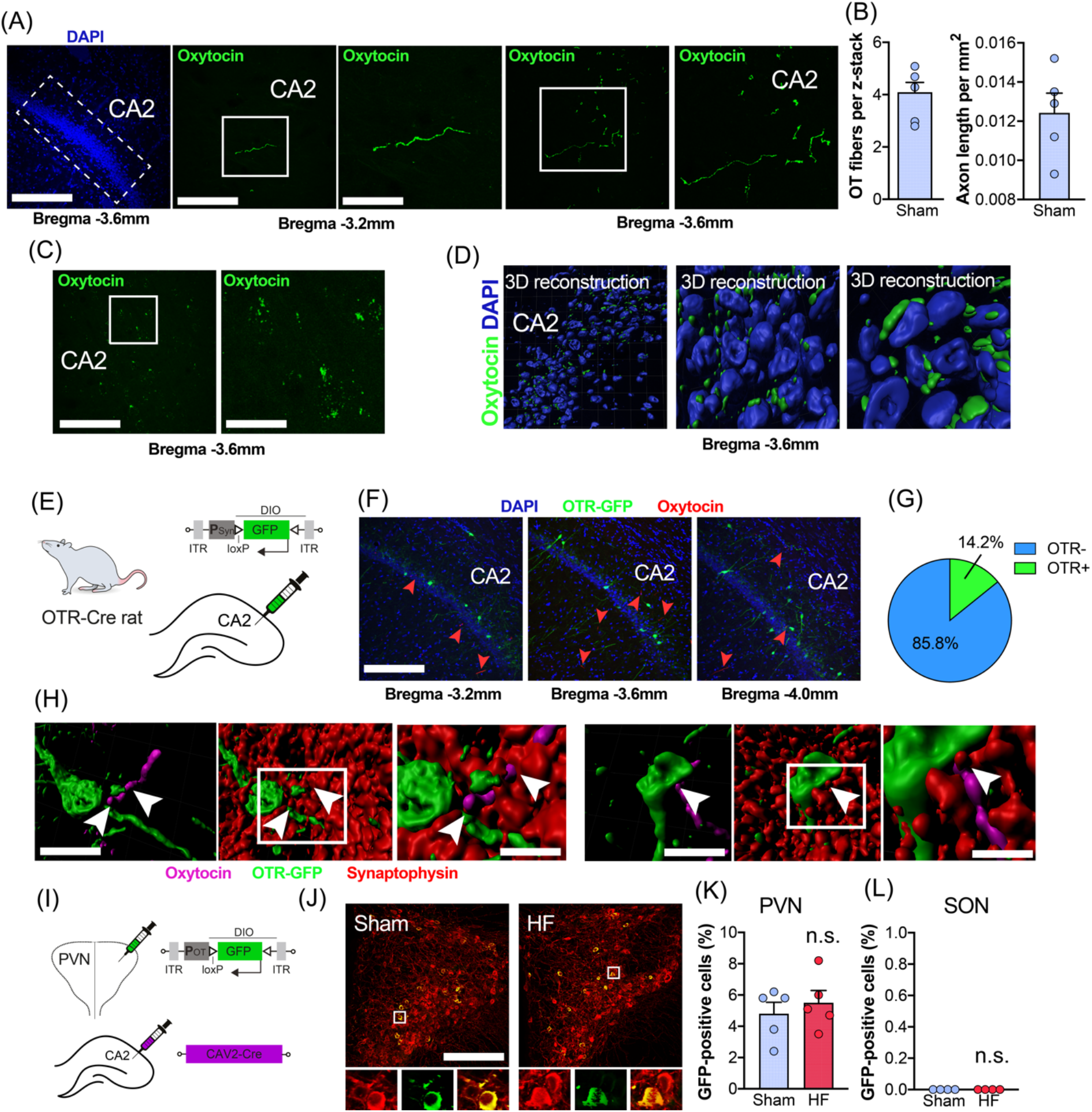
(A) Confocal images depict the presence of OTergic fibers in dorsal CA2. Scale bars = 250µm, 100µm and 50µm. Left image: dotted line indicates the location of dorsal CA2. Middle and right panels: white squares indicate regions of interest containing OTergic fibers, which are magnified in the right images, respectively. (B) Average number of OT-positive fibers per 40µm z-stack containing dorsal CA2 at Bregma −3.6mm. (C) OTergic fibers in CA2 form basket-like structures and were visualized via the OT (Neurophysin I) PS38 antibody. (D) Three-dimensional reconstructions of OTergic signal reveals co-localization of OT fibers with CA2 hippocampal neurons. OT-positive puncta appear in close proximity to CA2 neurons labeled with DAPI. Three-dimensional reconstructions show cells in the pyramidal cell layer. (E) Schematic depiction of the viral strategy to label OTR-positive neurons within hippocampal CA2 via injections of **P**Syn-DIO-GFP using the newly generated OTR-Cre rat line. (F) Confocal images show the location of OTR-positive cells (green, GFP) and OTergic fibers (red, arrowheads) within hippocampal CA2. Scale bar=250µm. (G) Pie chart shows the percentage of OTR-positive neurons (14.2%, n=3) in CA2. (H) Three-dimensional reconstruction of OTR-positive neurons (green), OTergic fibers (magenta) and the presynaptic marker synaptophysin (red) reveal potential direct OTergic innervation of OTR-positive cells in hippocampal CA2. Scale bar=25µm and 10µm. (I) Dual viral strategy for the retrograde labeling of CA2-projecting OT cells in SON and PVN. Rats were injected bilaterally with CAV2-Cre into hippocampal CA2 and bilaterally into either PVN or SON with **P**OT-DIO-GFP. (J) Confocal images show retrotraced, CA2-projection OT cells (yellow, green and red) in sham and HF rats. Scale bar=250µm. High magnification insets show overlap of OT immunoreactivity (red) and virally expressed GFP (green). Scale bar=20µm. (K) No difference in the number of retrotraced OT cells between sham and HF rats in the PVN (n=5 per group). (L) Injections of **P**OT-DIO-GFP into the SON of sham or HF rats did not result in labeling of OT neurons (n=5 per group).

OTR-expressing neurons in hippocampal CA2 have been reported in mice (16), but the exact location and percentage of OTR-positive cells in rat CA2 is – to the best of our knowledge – currently unknown. To this end, we employed the recently developed OTR-Ires-Cre knockin rats (27) (**Figure 2E**), which we injected with AAV-**P**Syn-DIO-GFP into CA2 to cell type-specifically express GFP in OTR-expressing neurons (n=3). We found that 14.2% of sampled neurons (**Figure 2F, G**, 409/2880 neurons, 8 sections per animal) of CA2 neurons were GFP-positive, some of which were in close proximity to OT-ergic immunoreactive fiber terminals (**Figure 2F, H**). This prompted us to probe for direct synaptic OTergic PVN→ CA2 contacts. We stained brain sections of injected OTR-Ires-Cre rats containing CA2 with PS38 for OT, and the presynaptic marker synaptophysin, and performed three-dimensional reconstruction of OTR-positive neurons (12), OTergic fibers and synaptophysin immunoreactivity (**Figure 2H**). As shown **in Figure 2H** (arrowheads) we occasionally observed direct, synaptophysin-positive, OTergic contacts (11/95 fibers, 11.6%) of soma and initial dendritic segments of OTR-positive neurons in CA2.

To further corroborate these findings, and to probe for potential differences in the number of CA2-projecting PVN OT neurons between sham and HF rats, we injected the monosynaptically-spreading CAV2-Cre virus into hippocampal CA2 and **P**OT-DIO-GFP into the ipsilateral PVN and SON (**Figure 2I**, n=5 per group). This approach allows viral-mediated fluorescent labeling of PVN OT neurons that synapse onto CA2 neurons. We found no difference in the number of retrogradely labeled, GFP-positive OT neurons in the PVN of sham and HF rats (**Figure 2J, K**), suggesting similar numbers of PVN→ CA2 projections. While we did not aim specifically to discriminate between magnocellular and parvocellular OT neurons, we observed that back-labeled OT neurons had large (>20µm) and round somata, typical of magnocellular neurons (9, 28, 29), a finding which is in line with the notion that magnocellular OT neurons innervate forebrain regions (9). In addition, we did not observe any retrogradely labeled cells in the SON (**Figure 2L**), suggesting that the PVN is the primary source of OTergic innervation to CA2 in rats (19).

### Reduced number of OT fibers, OT mRNA and protein in CA2 of HF rats

In order to determine if OTR expression in CA2 neurons was altered during HF, we performed immunohistochemical staining using a relatively new commercially available OTR antibody (AVR-013, Alomone labs). To validate the specificity of this antibody, we stained sections of WT and OTRko mice (30) (**Figure 3A**). We chose the central lateral amygdala (CeL) as our region of interest because it reliably demonstrates specific OTR ligand binding and the nucleus is anatomically well defined in mice and rats (10, 19, 31-33). We assessed OTR intensity and density for AVR-013 and found drastic differences for both parameters between WT and OTRko mice. While WT mice displayed strong OTR immunoreactivity and a higher density of OTR confined to the CeL, OTR immunoreactivity in OTRko mice was drastically diminished (**Figure 3B)**. Thus, we conclude that AVR-013 is a reliable and specific OTR antibody suited to study OTR levels in the rodent brain. In fact, this antibody has recently been used by other labs and validated using both peptide pre-incubation and knockdown approaches (34, 35).

**Figure 3.**
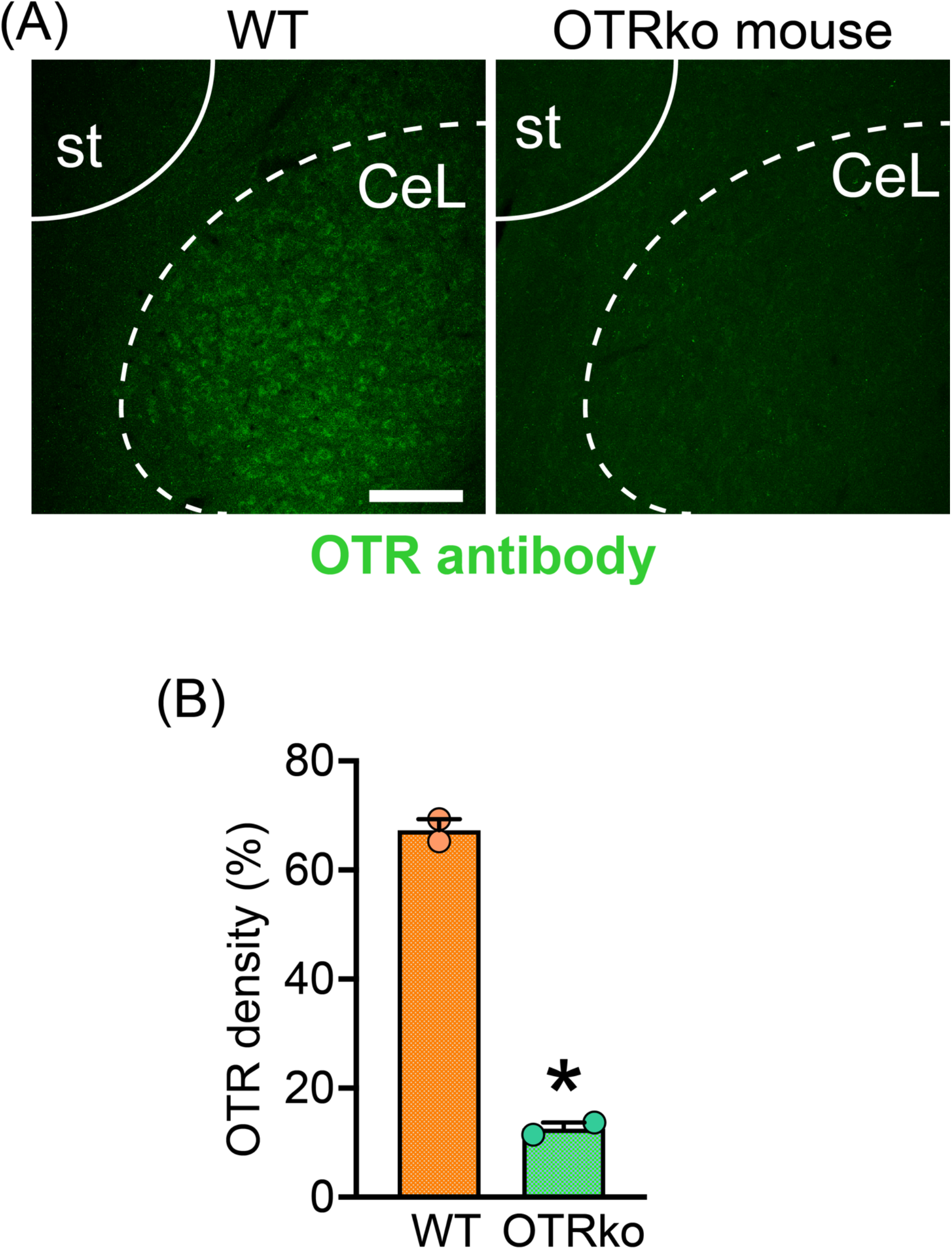
(A) Confocal images show brain sections containing the CeL of WT and OTRko mice stained with the AVR-013 OTR antibody. Scale bar=75µm. st=commissural stria terminalis, CeL=central lateral amygdala. (B) Decreased density of OTR immunoreactivity in the CeL of OTRko mice compared to WT mice. Student’s t-test, *p<0.05.

Next, we aimed to investigate potential HF-induced changes in PVN→ CA2 projections, as well as changes in OTR levels in the CA2 in this disease condition. To this end, we stained brain sections with the validated OTR antibody and analyzed OTR immunoreactivity in hippocampal CA2 of sham and HF rats (**Figure 4A, B**, n=5 per group). We found significantly diminished OTR immunoreactive intensity, as well density (area fraction) in HF rats, indicating an overall reduced expression of OTRs in this group (**Figure 4C**). Next, we analysed the number and intensity of immunoreactive OTergic projections in hippocampal CA2 of sham and HF rats (**Figure 4D**). We found significantly reduced OT signal intensity and density in HF rats, potentially indicating diminished OTergic innervation and/or depletion of OT content in PVN→ CA2 projections (**Figure 4E, F**). When we three-dimensionally reconstructed OTergic fibers within CA2, fibers in HF rats appeared thinner and seemed to contain less OT (**Figure 4G**). Next, we performed qPCR for OTR mRNA using tissue punches from PVN (21, 22) and CA2 (36) and found a 2-fold decrease in OTR mRNA in the PVN and a 3-fold decrease of OTR mRNA in CA2 of HF rats (**Figure 4H**). Finally, we performed Western blot for the OTR using brain region-specific punches of the CA2 and found significantly lower levels of OTRs in HF rats (**Figure 4I, J)**, thereby confirming our previous findings on OTR immunoreactivity (**Figure 4A-C**). Taken together, our results indicate significant changes to the OTergic PVN→ CA2 pathway, including diminished OTR mRNA and OTR protein in CA2, as well as a reduced number or content of OTergic fibers within the same structure.

**Figure 4.**
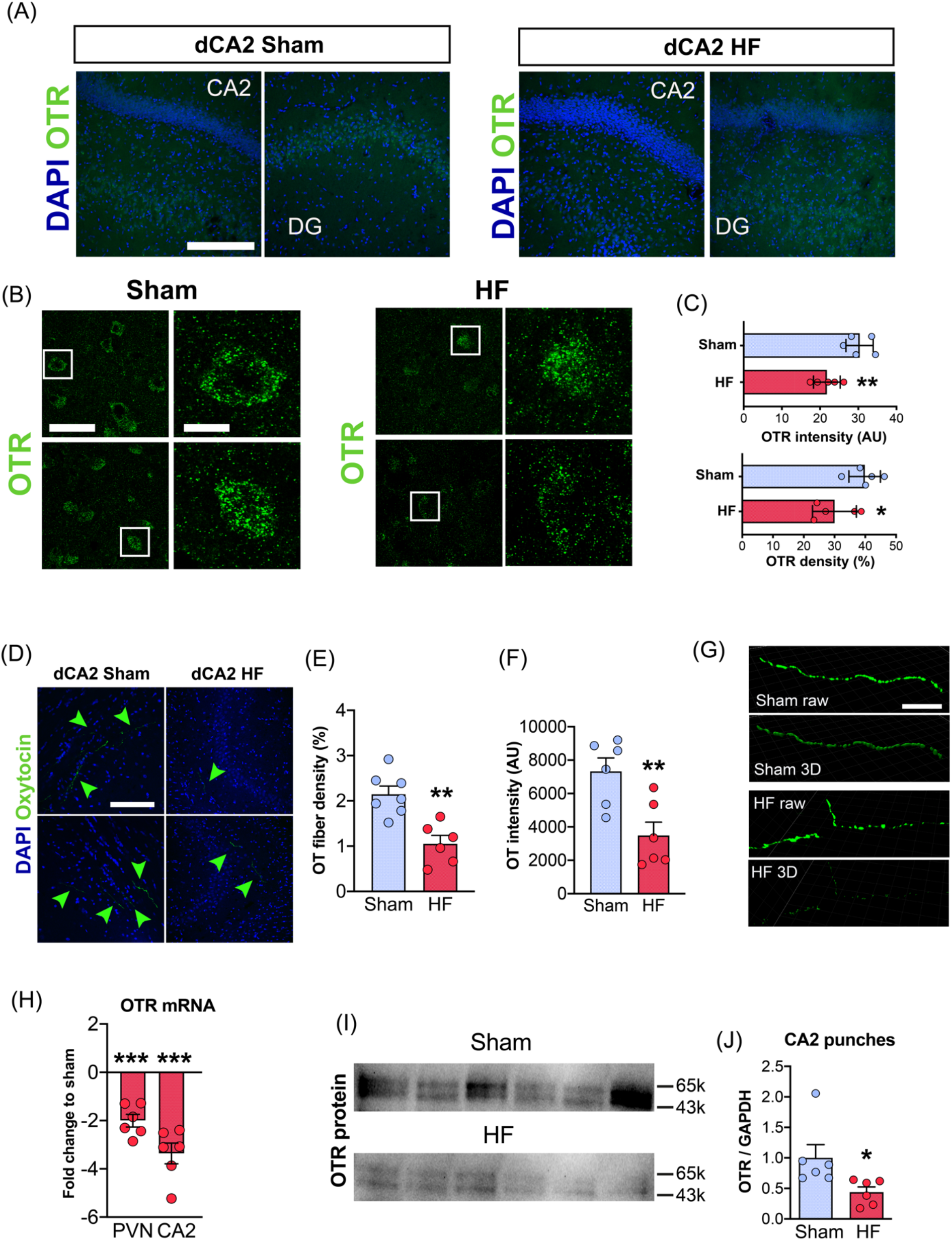
(A) Confocal images show dorsal hippocampus brain sections in a sham and a HF rat containing dentate gyrus (DG) and CA2 stained with an OTR antibody and labeled via DAPI. Scale bar=500µm. (B) High magnification image shows OTR immunoreactivity (green) in cells located in hippocampal CA2 for sham and HF rats. Scale bar=50 (left) and 15µm (right). (C) Reduced OTR intensity and density in CA2 if HF compared to Sham rats (n=5 per group). Data for OTR density and intensity were obtained using the entire confocal image. (D) Confocal images show OTergic fibers in hippocampal CA2 of sham and HF rats. Green arrowheads indicate location of OTergic (green) fibers. Scale bar=25µm. (E) Reduced OT fiber density in CA2 of HF rats (n=6 per group). (F) Reduced OT signal intensity in CA2 of HF rats (n=6 per group). (G) Raw and three-dimensionally reconstructed immunoreactive OTergic fibers of sham and HF rats in hippocampal CA2 visualized. Scale bar=5µm. (H) Decreased OTR mRNA levels in PVN and CA2 of HF rats assessed via tissue-specific punches and qPCR (n=6 per group). (I) Western blot images of OTR protein levels in CA2 of sham and HF rats. (J) Reduced OTR protein levels in CA2 of HF rats (n=6 per group). Student’s t-test or one-sample t-test, *p<0.05, **p<0.01 and ***p<0.0001.

### Diminished number of OTR-positive astrocytes in CA2 of HF rats

We recently described OTR-expressing astrocytes in the central amygdala of rats, which are crucial for the control and fine-tuning of the neuronal CeL→ CeM microcircuit (20). Thus, we were curious to determine whether other brain regions harbor OTR-expressing astrocytes, and thus performed double immunohistochemistry for the OTR and the astrocyte marker GFAP. We frequently observed OTR-positive astrocytes, identified by co-localization of GFAP and OTR immunoreactivity (47.7%, **Figure 5A**). When we compared sham and HF rats (n=3 per group), we found a stark reduction in the number of OTR-positive astrocytes (≈57% decrease) in HF rats (**Figure 5B, C**) (p< 0.0001 Chi-square test). To unequivocally demonstrate that hippocampal astrocytes are OTR-positive, we performed RNAScope in situ hybridization for OTR mRNA in combination with double immunohistochemistry against GFAP and glutamine synthetase (GS) (**Figure 5D**). Three-dimensional reconstruction of astrocytes and probe signal confirmed the presence of OTR mRNA inside of GS-labeled, GFAP-surrounded, astrocyte somata (red arrowheads, **Figure 5D**). We recently reported OTR-expressing astrocytes in the central amygdala, which are strategically-positioned to contact OTR-negative astrocytes, facilitate astrocyte-to-astrocyte communication through gap junctions, release D-serine in an activity-dependent manner and are morphologically distinct from OTR-negative astrocytes within the same nucleus (20). We thus analyzed whether OTR+ astrocytes in the CA2 display any distinct morphological features compared to OTR-negative astrocytes. Differently from what was previously described in the in the central nucleus of amygdala (20), we did not observe any distinct morphological parameters of OTR-positive, when compared to negative astrocytes (**Figure 5E**).

**Figure 5.**
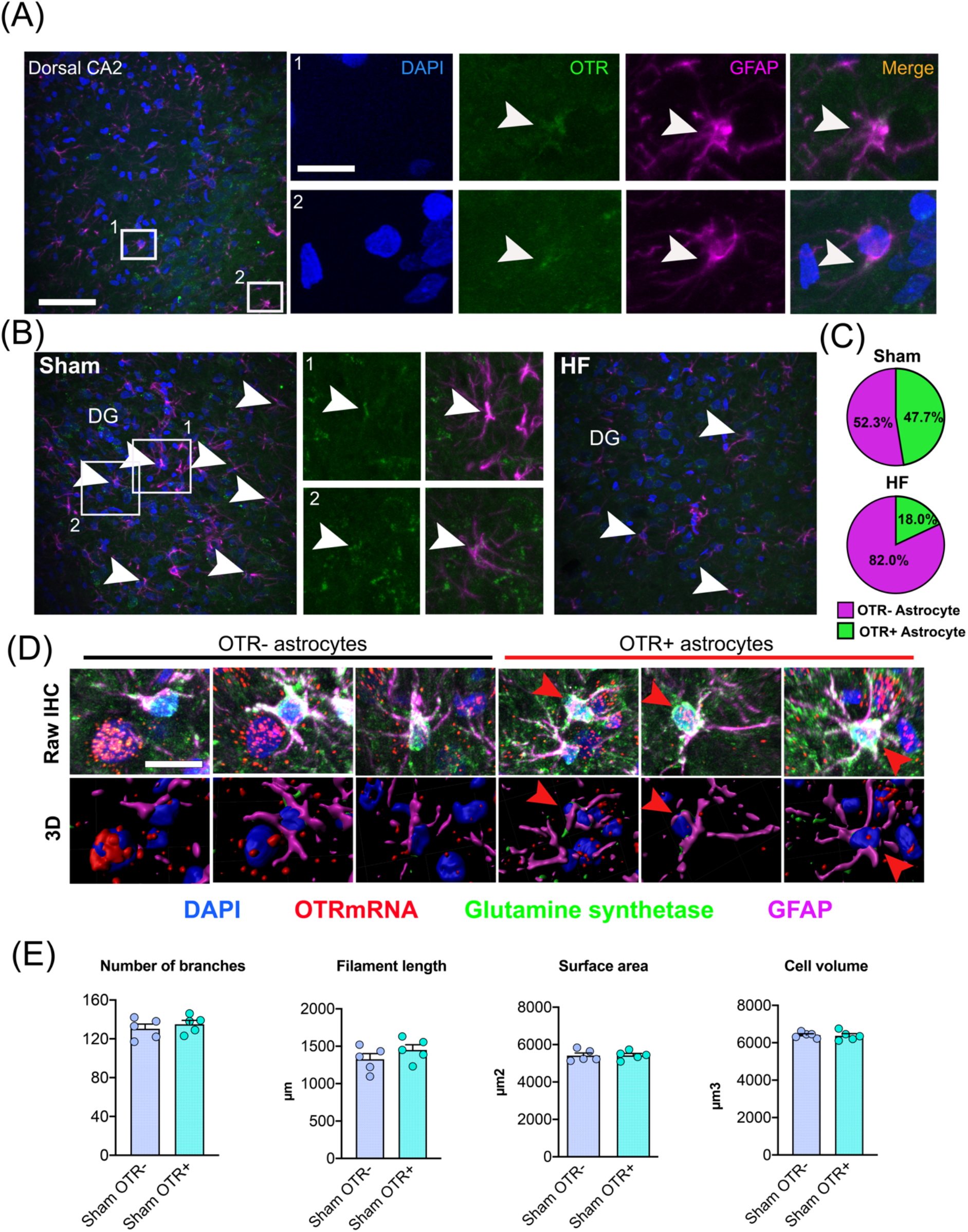
(A) Confocal images show the location and distribution of OTR-positive astrocytes in dorsal CA2 of a sham rat. Scale bar=100µm. High magnification insets show overlap of OTR immunoreactivity (green) and astrocyte processes (GFAP, magenta). White arrowheads indicate overlay of OTR and GFAP immunoreactivity. Scale bar=25µm. (B) Confocal images showing OTR-positive astrocytes in dorsal CA2 of a sham and a HF rat. White arrowheads indicate OTR-positive, GFAP-labeled astrocytes. DG=dentate gyrus. (C) Reduced number of OTR-positive astrocytes in HF rats (n=3 per group). (D) Raw immunohistochemical images and three-dimensional reconstructions of OTR-positive and OTR-negative astrocytes in dorsal CA2. Scale bar=15µm. (E) Three-dimensional astrocyte morphometric reveals no difference in the number of astrocyte branches, filament length, surface area or cell volume between OTR-negative and OTR-positive astrocytes in CA2 (n=5 per group).

Next, we compared sham and HF rats and found a significantly diminished number (≈50%) of OTR mRNA-positive astrocytes (**Figure 6A-C**), thus confirming our initial findings with the OTR antibody (**Figure 5B, C**). Intriguingly, when we analyzed the fraction of GS-negative, OTR mRNA-positive cells located in the pyramidal cell layer in CA2, we only observed a small reduction in HF rats, which did not reach statistical significance (**Figure 6D**, p=0.1135). Finally, we analyzed the number of astro-astrocytic contacts of OTR+/OTR-astrocytes with neighboring astrocytes in sham and HF rats using Imaris (20), and did not find any statistically significant differences between sham and HF rats (**Figure 6E**).

**Figure 6.**
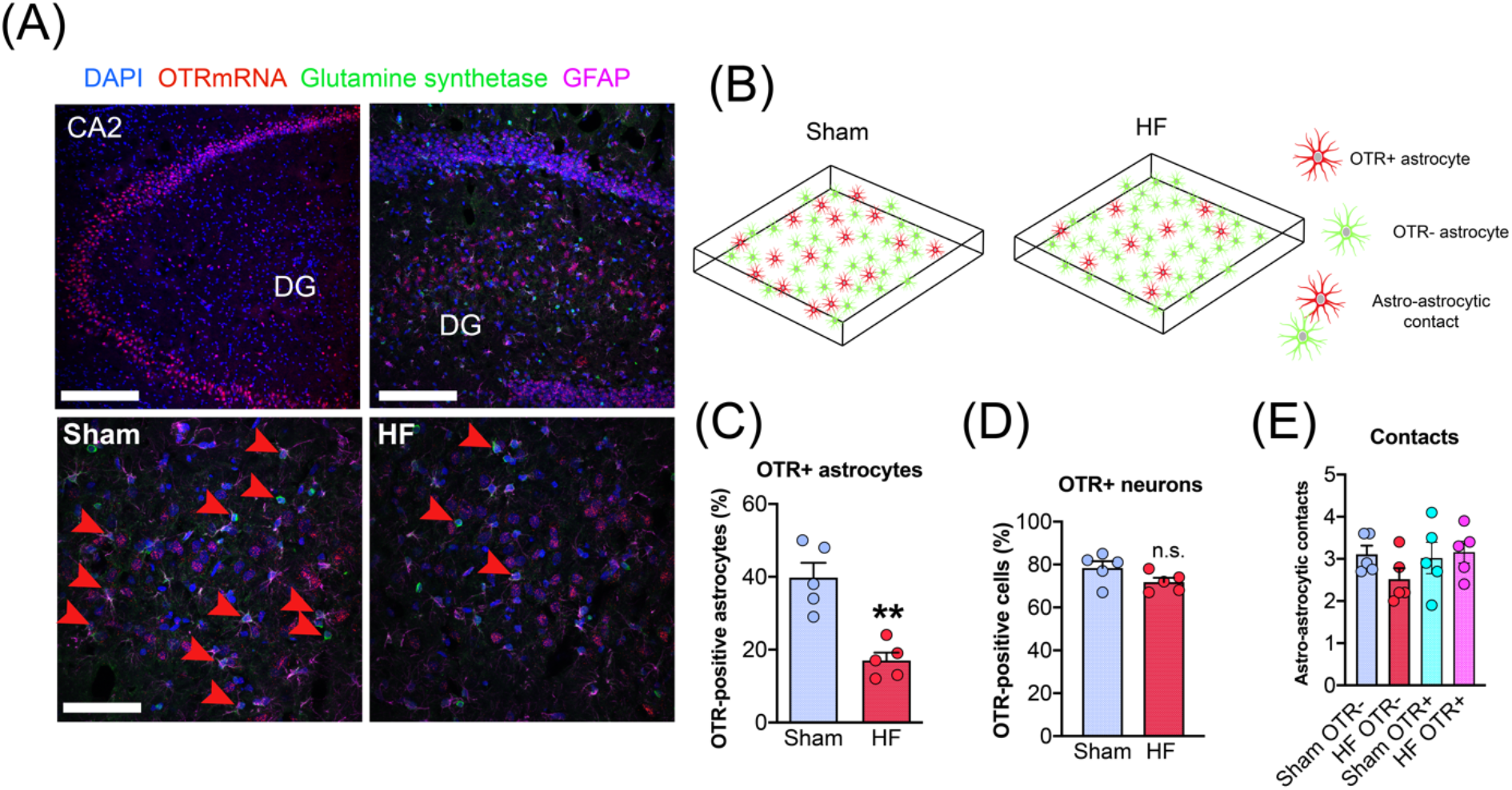
(A) Confocal images (top panels) show location and distribution of OTR mRNA-positive astrocytes labeled via glutamine synthetase and GFAP in the dorsal hippocampus. Scale bar=500µm, 250µm. Bottom panels show a reduced number of OTR-positive astrocytes in the CA2 of HF rats compared to sham. Red arrowheads indicate colocalization of GS immunoreactivity and OTR mRNA. Scale bar=100µm. (B) Schematic depiction of the random distribution of OTR-positive astrocytes in the CA2 of sham rats and reduced number of OTR-positive astrocytes in HF rats. (C) Decreased number of OTR-positive astrocytes in the CA2 of HF rats (n=5 per group). (D) No difference in the number of OTR-positive putative neurons (i.e. GS-negative cells, located in the pyramidal cell layers) between sham and HF rats. (E) No difference in the number of astro-astrocytic contacts between OTR-positive and OTR-negative astrocytes in sham and HF rats. Student’s t-test, **p<0.01.

## Discussion

Results from this study could be summarized as followed: I) The PVN (but not the SON) is a source of OTergic fibers to the hippocampal CA2 region; II) OT receptors (OTRs) within the CA2 are more abundantly expressed in astrocytes (∼40%) than in neurons (∼15%); III) In the PVN of HF rats we observed an increased OT immunoreactivity along with decreased OTR expression; IV) in the CA2 of HF rats, we observed both a blunted OTergic innervation, along with a diminished OTR expression, which appeared to be restricted to CA2 astrocytes.

### The role of hippocampal oxytocin signaling in health and disease

The hippocampus is among the most important brain regions for the acquisition, processing and storage of spatiotemporal, declarative and emotional memory (2, 4, 37). Impairment of proper hippocampal function due to neurodegenerative diseases such as Alzheimer’s disease (38), developmental complications (39) or damage (40) results in moderate to severe cognitive decline and memory loss in both rodents and humans (5, 6, 41-43). A role for the prosocial neuropeptide OT in the modulation of hippocampal memory formation, and its direct action on hippocampal neurons has been shown for different species and hippocampal subdivisions (14-16, 24, 44-47). For example, one study showed that an OT-driven hippocampal microcircuit involving the anterior dentate gyrus and anterior CA2/CA3 is crucial for the identification of social stimuli (46), and different authors showed that targeted deletion of OTRs within these regions resulted in impaired long-term social recognition in mice (44). Although OT shapes responses of principle cells of CA2 (16), likely facilitating social recognition, it remains enigmatic whether other cell types in CA2 express OTR and contribute to the OTergic modulation of the hippocampal microcircuit, especially in pathological conditions.

### The hypothalamic PVN is a source of OTergic innervation to CA2 in rats and mice

Previous publications reported OT fibers in the ventral hippocampus of mice (48), as well as in CA1, CA2 and CA3 of female rats (19, 49, 50). For example, Tirko and colleagues in a recent paper observed OTergic fibers on somata of hippocampal CA2 neurons in mice (16). In fact, the images shown on their Figure 1 for mice ((16), OT-YFP) look similar to what we occasionally observed in male Wistar rats (**Figure 2**). However, other studies could not reproduce those results (26). The contradictory nature of the findings could potentially be explained by a number of factors including differences in species or strains (i.e. mouse, Wistar rat or Sprague Dawley rat), sex (male/female), age (pups, adolescent or adult) and even individual variability (26).

A likely source of OTergic innervation of the hippocampus is the hypothalamic PVN, a region that provides OTergic innervation to multiple brain regions (19, 26, 51). Still, whether the PVN provides a direct OTergic innervation to the CA2 has also been controversial. In early studies, labs yielded opposing results so that OT fibers in the hippocampus either disappeared (52) or persisted (53) after targeted lesions of the PVN. In the present study we provide compelling evidence supporting the PVN as the primary, if not exclusive source of OTergic innervation to CA2 in rats – at least regarding hypothalamic OTergic sources. This is supported by our immunohistochemical studies showing synaptophysin-positive close appositions between OTergic fibers and OTR neurons in CA2. Furthermore, our retrograde tracing experiments via CAV2-Cre support more convincingly a direct contact of OTergic fibers originating in the PVN with neuronal somata in CA2 (**Figure 2**). A caveat however is that the CAV2-Cre virus can occasionally be taken up by axonal terminals in absence of direct contact with the postsynaptic neuron (54). Similar findings regarding OTergic innervation of CA2 has been described before in mice (16). Taken together, these results support direct projections of OTergic fibers to CA2 in both rats and mice, as well as synaptic-like contacts with OTR neurons in CA2.

### OTRs in hippocampal CA2 are predominantly expressed in astrocytes

For a thorough analysis of brain region- and cell type specific effects of OT under various conditions, it is paramount to have reliable tools for the assessment of variation in OTRs levels in the brain. While in situ hybridization techniques (20, 31, 49, 55) and autoradiography (56) are frequently used to visualize OTR mRNA or OTR protein, the use of OTR antibodies remains controversial (57). In fact, only certain batches of specific OTR antibodies seem to produce reliable results, which allow the quantification of OTR immunoreactivity (16, 58). Thus, we tested a relatively new OTR antibody (AVR-013, Alomone labs), which we validated using OTRko mice (**Figure 3**). We found that OTR immunoreactivity was essentially absent in OTR KO mice, thus confirming the specificity of the antibody. In fact, although this is a new batch of the OTR antibody, other labs have also successfully used the same (34, 35) or older batches (59), albeit without confirming its specificity via animal OTR KO models. We used this antibody to investigate neuronal and astrocytic OTR expression and obtained results (**Figure 4 and 5**) consistent with our data obtained via RNAScope (**Figure 5 and 6**), again confirming the efficacy and specificity of the antibody. We found that the number of OTR-positive neurons in the CA2 is relatively low (15%) and that the small number of direct OTergic contacts with OTR+ neurons as well as OTergic innervation is scarce (less than 4 fibers per 50µm, 10x confocal image). These findings seem to be in line with what has been reported previously in CA2 of mice (16). Following our previous demonstration of the existence of OTR+ astrocytes in the central amygdala of mice and rats (20) we wondered whether other brain regions might harbor OTR-expressing astrocytes as well. We chose CA2 as our region of interest due to the numerous reports of OTergic action within this region (15, 16, 44-46). However, to our big surprise, we found that in stark contrast to the number of OTR+ neurons (15%), almost half of the astrocytes located in CA2 expressed OTRs, as we obtained comparable numbers using either RNAScope for OTR mRNA (**Figure 6**) or the novel OTR antibody (**Figure 5**). Intriguingly, the proportion of OTR-positive astrocytes we observed in the hippocampus (39.8%) is higher than what we previously reported for the central amygdala (14-18%) (20). In addition, hippocampal OTR-positive astrocytes appeared not to be organized in a specific pattern as in the central nucleus of amygdala and were randomly positioned throughout the subdivisions of CA2 (**Figure 5**). Thus, it is tempting to propose that the mechanisms underlying OTR-mediated astrocyte activation in the hippocampus are fundamentally different from what we previously described in the central amygdala. This hypothesis needs to be validated with a series of astrocyte-specific recordings including a thorough analysis of the potential role of D-serine, which we pinpointed as the major gliotransmitter released from OTR-positive astrocytes in the central amygdala (20).

### Heart failure-induced changes to the PVN→ CA2 oxytocin pathway and OTR expression in astrocytes

In this study, we used the recently developed OTR-Cre rats (27) (27) to identify and analyze potential contacts of PVN OT fibers with OTR-positive neurons within the CA2. We report a series of HF-induced changes to the OT system both at the source and target of the PVN→CA2 OTergic pathway and hippocampus. In the PVN, we found an increase in OT immunoreactivity intensity, increased soma area, an overall increased OT cell volume and reduced OTR protein levels in the PVN of HF rats (**Figure 1**). These changes combined could reflect an increased activity of the PVN OTergic system during HF, leading to an enhanced local dendritic release of OT (60) and concomitant downregulation of OTR expression. This is consistent with earlier findings reporting overactivation of the OT system at early stages following a myocardial infarction (23). Conversely, we observed a reduction in the number of OTergic fibers in hippocampal CA2 with no concomitant changes in the number of retrotraced PVN OT neurons (**Figure 2**). These results suggest that the number of PVN→ CA2-projecting OT neurons is not diminished in HF rats. It is possible that PVN→ CA2-projecting OT neurons in HF rats form less bifurcations and weaker contacts in the target area, but given that we did not find any difference in the efficacy of CAV2-Cre-mediated retrolabeling of PVN OT neurons between sham and HF rats **(Figure 2**), we believe that this seems rather unlikely. Alternatively, these changes may reflect partial depletion of OT levels within the hippocampus due to an initial high activity levels of the PVN OTergic neurons, resulting in many fibers remaining below the detection threshold via IHC staining for OT. Future studies are needed to investigate the precise mode of OT release from those axons within CA2 and the precise time course of changes observed both in the PVN and CA2 during the course of HF.

Intriguingly, although we observed an overall reduction of OTR protein immunoreactivity (**Figure 4**), we only observed a significant reduction in OTR mRNA in astrocytes, but not neurons (**Figure 6**). Moreover, we observed a significant reduction in the total number of OTR-expressing astrocytes (**Figure 6C**), whereas the changes in OTR-positive neurons in HF rats did not reach statistical significance (**Figure 6D**). This seems particularly intriguing given that the role of OTRs in hippocampal astrocytes has not been studied yet, raising a series of important questions for future studies. For example, it is unclear whether OTergic release acts predominantly on astrocytes or neurons, and/or whether astrocytes act as initial gatekeepers, similar to what we recently described in the amygdala (20).

Taken together, our studies highlight astrocytes as key cellular targets mediating OTergic PVN inputs to the CA2 hippocampal region. Moreover, provides the first evidence for an altered PVN-to-CA2 OTergic system HF rats, which could potentially contribute to previously reported cognitive and mood impairments in this animal model.

### Perspectives

Critical questions arise from our study. For example, what mechanism in HF leads to selective downregulation of OTRs in astrocytes? It is known that HF initiates a plethora of short and long-term responses in the brain, including increased Angiotensin II signaling (61-63), hypoxia (64, 65) and neuroinflammation (22, 61). Thus, it is plausible that any (or several) of these mechanisms contribute to the downregulation of OTRs in hippocampal astrocytes. It seems particularly intriguing that we observed downregulation of OTRs in astrocytes, but not neurons. We recently reported elevated hypoxia markers Hif1α and Hif2α, as well as pathological vascularization of the hippocampus (36). Thus, it is possible that astrocytes are more susceptible to this hypoxic state. Finally, and perhaps more importantly, what are the potential behavioral consequences of the altered OTergic PVN→ CA2 pathway in HF? One could speculate that given the importance of hippocampal OT signaling in neuromodulation (14, 16, 49) and social memory (44, 46), pathological alterations of hippocampal networks might occur that result in cognitive impairments and memory loss like we recently reported (21).

Another intriguing aspect that arose from our study relates to the modality of OT release and action within the hippocampus. Given (1) the low number of OTR-positive neurons in the CA2; (2) the overall scarce OTergic innervation in this region (less than 4 fibers per 50µm, 10x confocal image; (3) the small number of direct OTergic contacts with OTR+ neurons; and (4) the relatively high number of OTR-expressing astrocytes, it is likely that a local “micro-volume” transmission modality is the predominant manner of OTergic release and action within CA2. These findings give rise to the exciting possibility that hippocampal OTergic release is not uniquely targeting neurons, but rather (or even predominantly) astrocytes as well.

Finally, given the recently identified role of OTR-positive astrocytes in the central amygdala in the modulation of pain and anxiety (20), it will be paramount to understand whether and how functions changes in OTR+ astrocytes contribute to neuropsychiatric and neurodegenerative diseases.

## Acknowledgements

We thank Lara Barteczko for packaging Cre-dependent rAAV.

## Data availability

All data generated in this study will be available from the corresponding author upon request.

